# Contribution of the novel oxazolidinone TBD09 (MK-7762) to regimens with bedaquiline and pretomanid in a mouse model of tuberculosis

**DOI:** 10.64898/2026.09.23.753971

**Authors:** Deepak V. Almeida, Paul J. Converse, Courtney Schill, Jin I. Lee, Micha Levi, Alexander Berg, Khisimuzi Mdluli, Eric L. Nuermberger

## Abstract

Linezolid is a key component of the bedaquiline, pretomanid, linezolid (BPaL) regimen for rifampin-resistant tuberculosis, but dose-and duration-dependent mitochondrial toxicity frequently limits its use. TBD09 (formerly MK-7762) is a novel oxazolidinone developed to maintain antitubercular potency comparable to linezolid while reducing mitochondrial protein synthesis inhibition and related adverse events. We evaluated the contribution of TBD09 to the bactericidal and sterilizing activities of BPa-based regimens in a murine tuberculosis model. BALB/c mice were aerosol-infected with *Mycobacterium tuberculosis* H37Rv and treated starting 2 weeks post-infection. In Experiment 1, dose-ranging bactericidal activity of TBD09 (50, 100, 200 mg/kg) was assessed alone and in combination with BPa, with comparison to linezolid and sutezolid. In Experiment 2, the sterilizing activity of TBD09 (200 mg/kg) in combination with BPa and BPa plus moxifloxacin (M) was assessed by evaluating relapse after treatment durations of 4-22 weeks. TBD09 monotherapy was bacteriostatic, similar to linezolid over the first 4 weeks of treatment. The addition of TBD09 to BPa significantly enhanced bactericidal activity comparable to the addition of linezolid. In relapse assessments, BPa+TBD09 resulted in significantly fewer relapses compared to BPa alone after 14 weeks of treatment, and BPaM+TBD09 resulted in significantly fewer relapses compared to BPaM after 10 weeks. Model-based analysis estimated the treatment duration to 95% cure for BPa+TBD09 at 2.7 weeks, compared to 3.2 weeks for BPaL, although the time to 50% cure was similar. These results demonstrate that TBD09 provides bactericidal and sterilizing activity similar to linezolid in combination with BPa(M), supporting further clinical development of TBD09 for tuberculosis.

## INTRODUCTION

Tuberculosis (TB), caused by *Mycobacterium tuberculosis*, continues to be the leading infectious cause of mortality worldwide. Most people with TB can be treated successfully with the standard four-drug regimen of rifampin, isoniazid, ethambutol, and pyrazinamide. Rifampin-resistant (RR-) TB requires treatment with alternative regimens, including the recently recommended regimen of bedaquiline (B), pretomanid (Pa), and linezolid (L), with or without moxifloxacin (M)(1), which represents the first oral short-course (6-month) regimen for RR-TB. However, prolonged use of linezolid can be associated with potentially severe neurological and hematological side effects that are dose-and duration-dependent. As a result, a more selective alternative oxazolidinone antibiotic that retains efficacy comparable to linezolid without these side effects is highly desirable.

TBD09 (MK-7762) is a novel oxazolidinone developed to maintain antitubercular potency comparable to linezolid while reducing the potency of human mitochondrial protein synthesis inhibition to reduce the risk of related adverse events, such as anemia and neuropathy (2). It has demonstrated an improved preclinical safety profile and may be more suitable for once-daily dosing than linezolid (2). The safety, tolerability and pharmacokinetics of TBD09 are now being evaluated in Phase 1 trials.

Although the dose-ranging bactericidal activity of TBD09 monotherapy in acute and chronic infection models in BALB/c mice was reported recently (2), it remains to be determined if TBD09 can be substituted for linezolid in the BPaL and BPaLM combinations without loss of efficacy and how it may compare to other oxazolidinones in this regard. Using a subacute mouse infection model in which the promising sterilizing activity of the BPaL(M) regimen was first shown (3, 4), we carried out a dose-ranging experiment to evaluate the bactericidal activity of TBD09 as monotherapy and in combination with BPa, comparing its activity to that of linezolid and sutezolid (U), another oxazolidinone in clinical development. Confirming that the addition of TBD09 increased the bactericidal activity of the BPa regimen backbone, we proceeded to assess the sterilizing activity of TBD09 in combination with BPa and BPaM, with comparison to sutezolid. The results confirm the bactericidal and sterilizing activity of TBD09 in this murine model and support its further clinical development.

## RESULTS

### Minimum inhibitory concentration (MIC)

To compare the antitubercular potency of TBD09, linezolid, and sutezolid, their respective MICs were determined against *M. tuberculosis* H37Rv using the broth macrodilution method in 7H9 medium. The MIC of TBD09 was 2.0 µg/ml versus 1.0 µg/ml for linezolid and 0.25 µg/ml for sutezolid. Mean MICs of TBD09 and linezolid were also determined using a resazurin microplate assay and found to be 1.6 and 1.2 µg/ml, respectively.

### Experiment 1: Dose-ranging bactericidal activity of TBD09, alone and in combination with bedaquiline and pretomanid, in a subacute mouse model of TB

Experiment 1 evaluated the dose-ranging activity of TBD09 when given at doses of 50, 100 and 200 mg/kg, both alone and in combination with the BPa backbone, with comparisons to linezolid 100 mg/kg and sutezolid 50 mg/kg after 4 and 8 weeks of treatment. The scheme of the experiment is presented in **Table S1**. BALB/c mice were infected via aerosol with approximately 4 log_10_ CFU of *M. tuberculosis* H37Rv. Two weeks later, at the start of treatment (D0), the mean CFU count in the lungs was 7.63±0.06. As shown in **FIG 1** and **Table S2**, regardless of TBD09 dose of 50, 100, or 200 mg/kg, mean lung CFU counts after 4 weeks of treatment (range 7.44 to 7.65 log_10_) were similar to those observed with linezolid (7.52 log_10_) and indicated bacteriostatic activity. In contrast, sutezolid monotherapy was bactericidal (6.66 log_10_; p<0.0001) compared to Day 0 and more active than TBD09 at all doses or linezolid alone at this time point (all, p<0.0001). BPa was bactericidal, resulting in a mean lung CFU count of 6.47 log_10_, and this activity was significantly enhanced (p<0.0001) by the addition of each oxazolidinone. As with the monotherapy arms, no dose-response relationship was observed for TBD09 in combination with BPa and its additive effect was similar to that of linezolid, whereas the addition of sutezolid enhanced the activity of BPa to a greater extent. After 8 weeks of treatment, mean lung CFU counts were modestly lower than the baseline count in mice treated with TBD09, ranging from 7.06 to 7.25 log_10_, whereas both linezolid and sutezolid were more bactericidal, resulting in mean lung CFU counts of 6.28 and 5.11, respectively. Although no dose-response relationship for TBD09 was observed in monotherapy, the bactericidal activity of TBD09 did increase with dose in combination with BPa. The combination of BPa+TBD09 at 200 mg/kg was more active (p=0.0322) than BPa with linezolid at 100 mg/kg but not as active as BPaU (p=0.0498). The activity of BPa+TBD09 at 50 and 100 mg/kg was not significantly different (p=0.0561 and 0.5159, respectively) compared to that of BPa with linezolid at 100 mg/kg.

**FIG 1.**
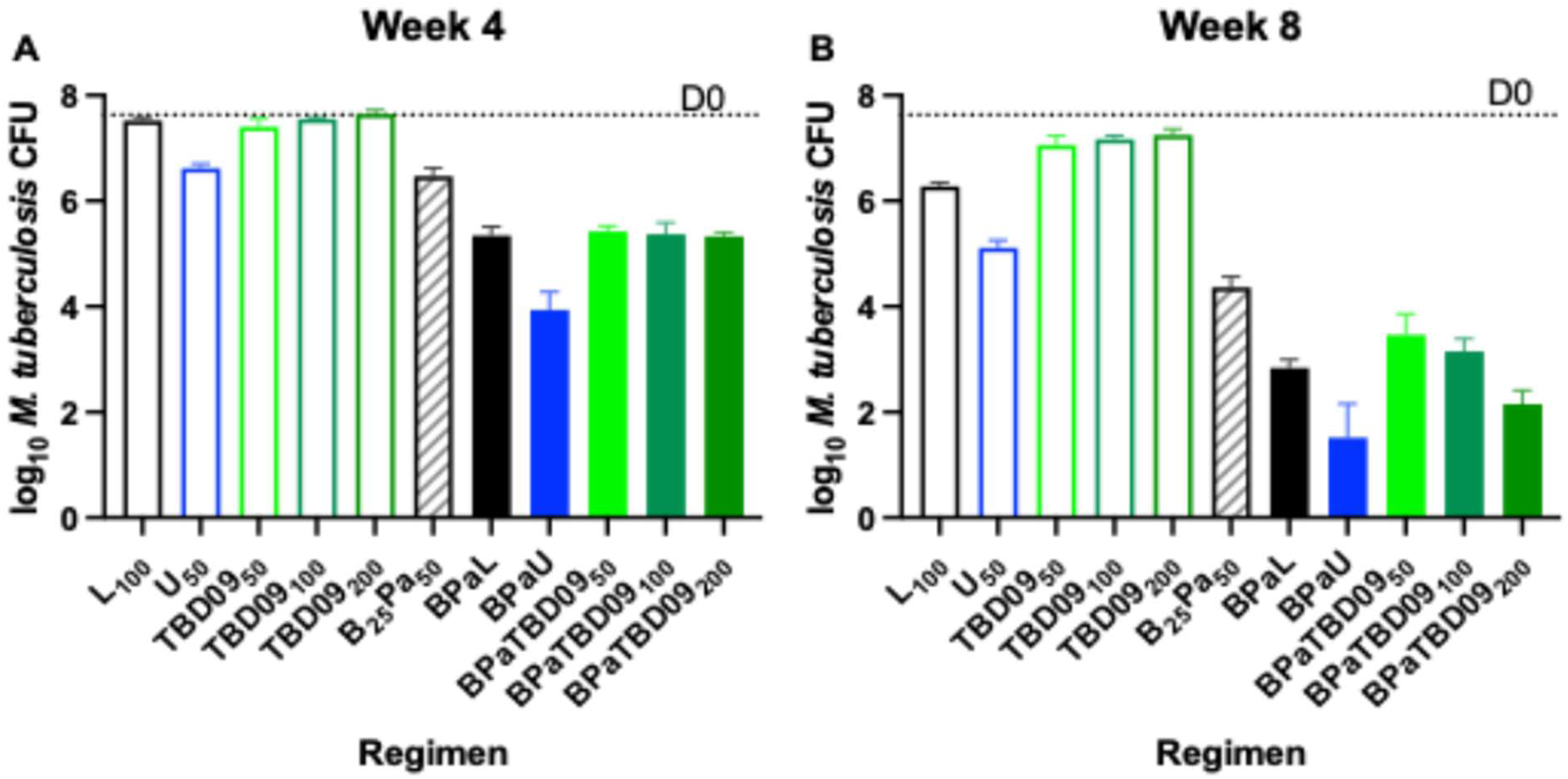
Mean lung CFU results after 4 and 8 weeks of treatment in Experiment 1. Oxazolidinones as monotherapy in open bars and in combination with BPa in solid bars.

### Pharmacokinetic analysis

Pharmacokinetic (PK) analysis was performed to determine the drug exposure and to compare drug exposures in infected mice receiving TBD09 alone and in combination with BPa, as well as BPa alone. Sparse sampling to enable measurement of TBD09, bedaquiline and pretomanid whole blood concentrations was performed during the second week of treatment. TBD09 showed dose-dependent exposures that did not differ when TBD09 was administered alone or in combination with BPa. Drug concentrations in the BPa+TBD09_200_ arm could not be determined due to technical issues. For all other groups, the whole blood concentrations of TBD09 increased in a less-than-dose-proportional manner (**Table 1**) but exceeded the MIC of 2 µg/mL for all or most of the dosing interval. Concentrations of bedaquiline and its M2 metabolite, and those of pretomanid, were not affected by co-administration with different doses of TBD09 (**Table S3** and **FIG S1**).

**Table 1.**
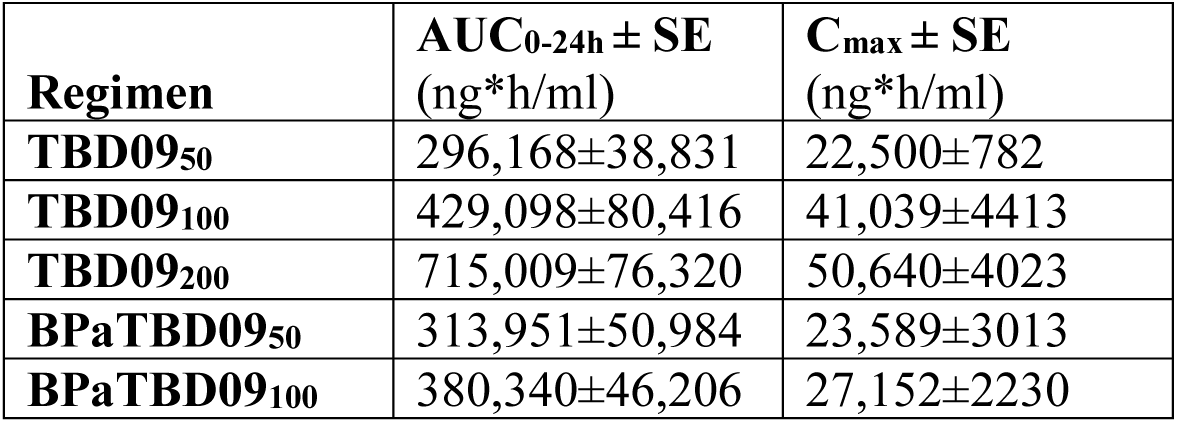
Mean plasma pharmacokinetic parameters for TBD09 in Experiment 1.

### Experiment 2: Bactericidal and sterilizing activity of TBD09 in combination with BPa or BPaM in a relapsing mouse model of TB

Experiment 1 demonstrated that addition of TBD09 to BPa increased the bactericidal activity of the combination in dose-dependent fashion. Experiment 2 evaluated the contribution of TBD09 at the highest dose tested in Experiment 1, 200 mg/kg, to the bactericidal and sterilizing activity of the BPa and BPaM regimen backbones, with comparison to that of sutezolid. BPaL and BPaM plus pyrazinamide (i.e., BPaMZ) were included as clinical comparators. The scheme of Experiment 2 is presented in **Table S4**. In the first week of treatment, one mouse each from the BPaL, BPaM and BPaMU arms died due to gavage-related injury. After two weeks of treatment, the addition of linezolid, sutezolid, TBD09, and moxifloxacin each significantly increased (p=0.0083, p<0.0001, p=0.0085, and p=0.0342, respectively) the bactericidal activity of the BPa backbone compared to BPa alone (**FIG 2 and Table S5**). The mean lung CFU count in mice treated with BPa+TBD09 was not significantly different from that in mice receiving BPaL but mice treated with BPaU had a significantly (p<0.0001) lower mean CFU count than mice treated with BPaL. Compared to treatment with BPaM, treatment with BPaM plus a fourth drug (i.e., TBD09, sutezolid, or pyrazinamide) resulted in significantly (p=0.0011, p<0.0001, and p<0.0001, respectively) lower CFU counts. BPaMZ was not significantly more bactericidal than BPaMU, but it was more active (p=0.0003) than BPaM+TBD09.

**FIG 2.**
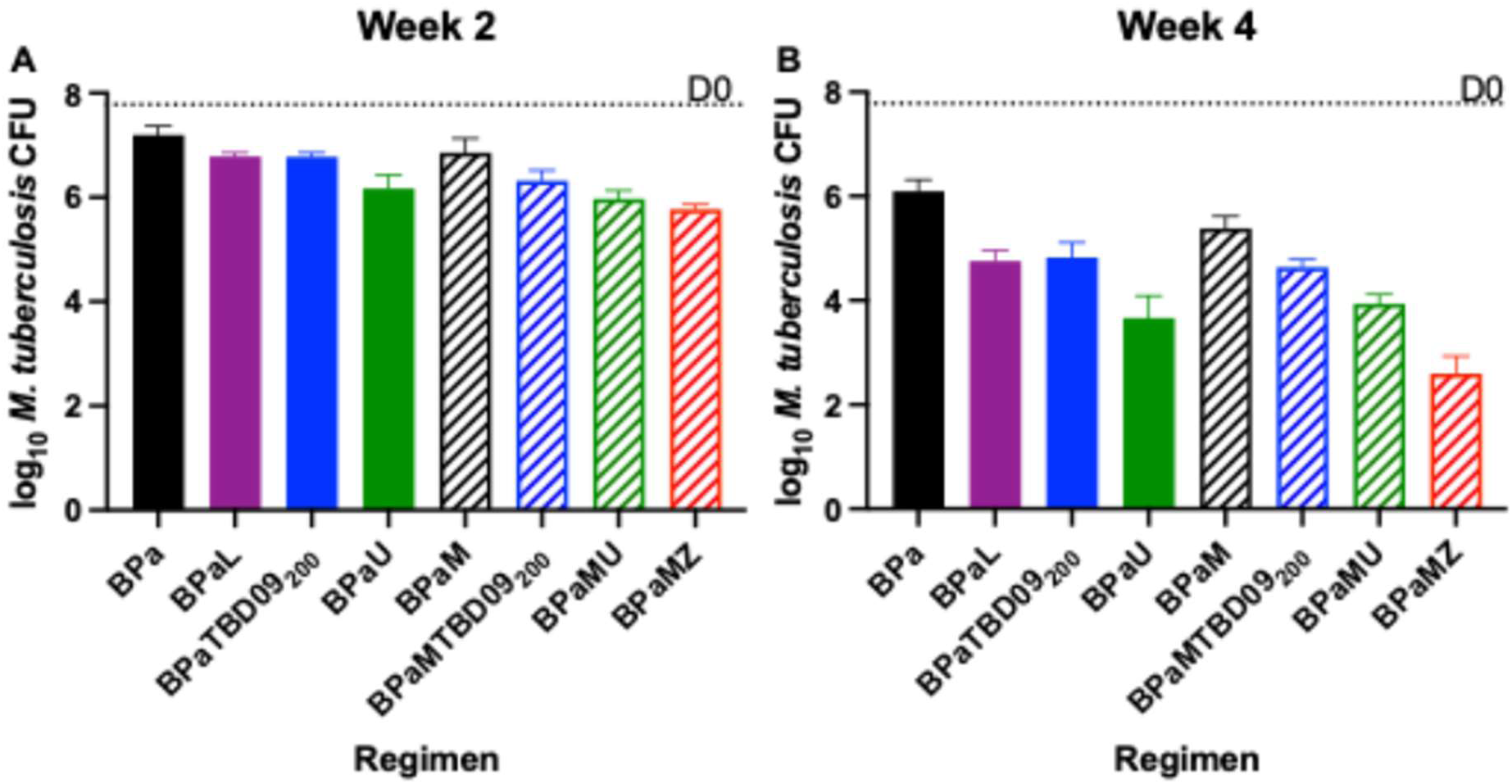
Mean lung CFU results after 2 and 4 weeks of treatment in Experiment 2. BPa and BPa + oxazolidinone are shown in solid bars and BPaM-based regimens are shown in diagonally striped bars.

After 4 weeks of treatment, all 3-drug regimens again resulted in significantly lower mean CFU counts compared to BPa (p<0.0001 for BPaL, BPa+TBD09, and BPaU; and p=0.0029 for BPaM). Compared to BPaL, BPaU treatment resulted in significantly lower CFU counts (p=0.0001) but BPa+TBD09 was not significantly different, as observed at this time point in Experiment 1. Compared to BPaM, all 4-drug regimens resulted in significantly lower CFU counts (p=0.0005 for BPaM+TBD09 and p<0.0001 for BPaMU and BPaMZ). BPaMZ was significantly (p<0.0001) more bactericidal than BPaMU, which was significantly more bactericidal than BPaM+TBD09.

The contribution of TBD09 to the sterilizing activity of BPa and BPaM was assessed by evaluating for relapse in cohorts of 3-5 mice treated for durations ranging between 4 and 22 weeks before being left untreated for an additional 12 weeks. As shown in Table 2, only 1 of 5 mice treated with BPaMZ relapsed after 6 and 8 weeks of treatment and all were relapse-free after 10 weeks of treatment, similar to previously published results (4).

**Table 2.** Proportions of mice relapsing after treatment with the indicated regimens in Experiment 2.

| Treatment | Time point and proportion of mice relapsing |  |  |  |  |  |  |  |  |
| --- | --- | --- | --- | --- | --- | --- | --- | --- | --- |
|  | W4<br>(+12) | W6<br>(+12) | W8<br>(+12) | W10<br>(+12) | W12<br>(+12) | W14<br>(+12) | W16<br>(+12) | W19<br>(+12) | W22<br>(+12) |
| B<sub>25</sub>Pa<sub>50</sub> |  |  |  | 4/4<sup>1</sup> | 5/5 | 5/5 | 3/5 | 0/4 | 0/3 |
| BP<sub>a</sub>U<sub>50</sub> |  | 3/4 | 2/5 | 0/5 | 3/5 | 0/4 | 0/3 |  |  |
| BP<sub>a</sub>L<sub>100</sub> |  | 3/3<sup>2</sup> | 5/5 | 1/5 | 1/5 | 0/5 | 0/3 | 0/3 |  |
| BP<sub>a</sub>TBD09<sub>200</sub> |  | 4/4 | 5/5 | 4/5 | 2/5 | 1/5 | 0/3 | 0/3 |  |
| BP<sub>a</sub>M<sub>100</sub> | 3/3<sup>2</sup> |  | 5/5 | 5/5 | 3/5 | 1/5 | 0/3 | 0/3 |  |
| 8wBP<sub>a</sub>MZ<sub>150</sub>/<br>BP<sub>a</sub>M | 4/4 | 1/5 | 1/5 | 0/4 |  |  |  |  |  |
| BP<sub>a</sub>MU | 3/3 | 4/4<sup>2</sup> | 3/5 | 1/5 | 2/5 | 0/3 |  |  |  |
| BP<sub>a</sub>MTBD09 |  | 5/5 | 5/5 | 0/5 | 2/5 | 0/5 | 0/5 |  |  |
| <sup>1</sup>Includes one mouse counted as relapse after being euthanized for neck abscess at W10 (+5) and found to have detectable CFU in lung homogenate. <sup>2</sup>One mouse in each of these groups died due to a gavage accident during the first week of treatment and was not assessed for the relapse endpoint. |  |  |  |  |  |  |  |  |  |

BPaL required an additional 4 weeks of treatment to achieve similar results. Compared to mice treated with BPa alone, those treated with BPa+TBD09 were more likely to be relapse-free after 10-16 weeks of treatment; and this difference was statistically significant after 14 weeks of treatment (p=0.0476), despite the fact that this pairwise analysis was underpowered. Compared to mice treated with BPaM alone, those treated with BPaM+TBD09 had a significantly (p=0.0079) lower proportion with relapse after 10 weeks of treatment. Although TBD09 and sutezolid contributed similar sterilizing activity to the BPaM backbone, these combinations were not as rapidly sterilizing as BPaMZ. Model-based estimation of the probability of cure (i.e., no relapse) according to treatment duration for each regimen is shown in **FIG 3**, with test regimens comprised of BPa plus an oxazolidinone shown in **FIG 3A** and test regimens containing BPaM shown in **FIG 3B**. The estimated treatment duration to 95% cure (T95) values, equivalent to treatment duration resulting in a 5% relapse probability, are shown in **Table S6**. Compared to BPaL, BPaTBD09 had a lower T95 value but a similar T50 value and did not have fewer observed relapses at any time point, making it difficult to differentiate the activity of these two regimens. For BPaU, the confidence interval around the T95 estimate was very large due to the surprising observation of 3 relapses at W12(+12), which was higher than the number of relapses observed at W8(+12) and W10(+12) combined. One of the W12(+12) relapses was attributable to detection of a single CFU within the entire lung homogenate. Note that model-based estimates were generated by pooling the study data with historical studies and accounted for inter-study variability and covariate effects, specifically the *M. tuberculosis* inoculum amount used in the study (5). After adjustment for these effects, the model-estimated curves show slightly different profiles than the pair-wise comparisons at specific timepoints (**FIG S2**).

**FIG 3.**
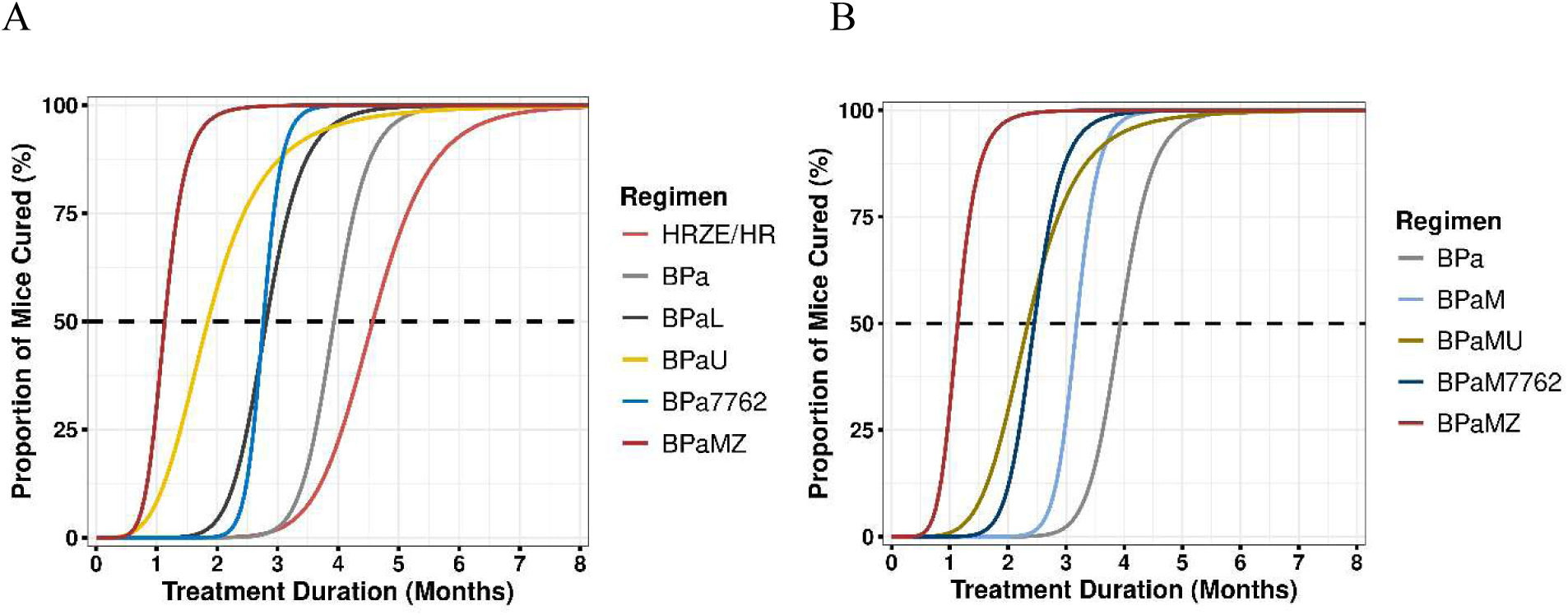
Statistical modelling curves depicting the proportion of mice cured (i.e., relapse-free) by duration of treatment with test regimens comprised of adding oxazolidinones to the (A) BPa backbone or (B) BPaM backbone

## DISCUSSION

Treatment of tuberculosis, especially RR-TB, is challenging due to the limited number of effective and tolerable drugs (6). The introduction and implementation of the BPaL(M) regimen enabled effective treatment of RR-TB with shorter durations of oral drug therapy. However, linezolid causes dose-and duration-dependent mitochondrial toxicity in the form of peripheral blood cytopenias, peripheral and optic neuropathies, and lactic acidosis (7). These adverse effects require clinical and laboratory monitoring and are often treatment-limiting. In cases of peripheral and optic neuropathy, adverse effects and disability can be irreversible (8). Replacement of linezolid with a safer and similarly efficacious oxazolidinone would significantly improve the implementation of safe and effective short-course RR-TB treatment and may also enable BPa-based regimens to be used more widely as alternatives to rifamycin-based regimens for drug-susceptible TB.

TBD09 is an orally available oxazolidinone developed for greater selectivity against mycobacterial ribosomes compared to linezolid (2). It demonstrated a favorable preclinical safety profile and recently completed phase 1 clinical trials in which it was well-tolerated with no safety concerns identified; and once-daily doses produced exposures exceeding the MIC for the entire dosing interval (2, 9). Specifically, TBD09 doses achieving average plasma trough concentrations approximately 5-18 times above MIC did not adversely affect hematological safety laboratory parameters over 28 days of administration (9). In this context, our finding that TBD09 adds bactericidal activity and sterilizing activity to the BPa backbone is very encouraging. At a dose of 100 mg/kg, TBD09 added bactericidal activity indistinguishable from the same dose of linezolid in Experiment 1. At the only dose tested in Experiment 2, 200 mg/kg, TBD09 added significant sterilizing activity to BPa that may not have been as great as that of linezolid at 100 mg/kg in terms of the absolute number of relapses enumerated but was estimated to be similar to linezolid in our model-based analysis that takes prior performance of BPa and BPaL into account. Although neither linezolid nor TBD09 was as potent as sutezolid in combination with BPa, they were each superior to moxifloxacin in this model.

Although the most predictive PK/PD parameter for oxazolidinone activity against *M. tuberculosis* remains uncertain, especially in combination therapy, the average unbound plasma AUC/MIC in our model was generally similar for linezolid at 100 mg/kg (i.e., approximately 171.5 µg-h/ml) and TBD09 at 100-200 mg/kg (i.e., approximately 149-270 µg-h/ml), which aligns well with their potency observed here in combination with BPa in mice. Against a panel of clinical *M. tuberculosis* isolates and reference strains, TBD09 MICs were the same or lower than linezolid MICs (i.e., MIC_90_ of TBD09 = 0.78 µM (0.33 µg-h/ml) vs. 1.56 µM for linezolid) (2). Therefore, TBD09 may have at least similar activity to linezolid if similar exposures (i.e., AUCs) are achieved. In the recently completed phase 1 trial, steady state plasma AUC_0-24h_ values of approximately 169 and 279 µg-h/ml were observed in participants receiving daily TBD09 doses of 300 mg and 500 mg, respectively, with food (9), comparable to previously published AUC ranges observed with 600-1200 mg total daily doses of linezolid (10, 11). Thus, TBD09 at these doses may safely achieve AUCs similar to 600-1200 mg of linezolid, while also maintaining TBD09 concentrations several fold above MIC for the entire dosing interval given its longer half-life.

Remarkably, when administered as monotherapy in our subacute high-dose aerosol mouse infection model, TBD09 was at least as potent (based on mg/kg dose) as linezolid over the first 4 weeks of treatment but was less potent and less effective than linezolid over 8 weeks of treatment. Moreover, no dose-response relationship was observed for TBD09, suggesting a lower maximal effect of TBD09 compared to linezolid. This apparent difference in pharmacodynamics of TBD09 and linezolid was limited to the condition of monotherapy, as TBD09 exhibited dose-dependent additive bactericidal activity that was at least as potent as linezolid when combined with BPa for 8 weeks. The lower maximal effect of TBD09 monotherapy compared to linezolid monotherapy at 8 weeks remains unexplained. Acquired drug resistance through selection of spontaneous drug-resistant mutants is an unlikely explanation. Mutations in *rrl* and *rplC* associated with oxazolidinone resistance are very rare amongst wild type populations (12). Crowley et al (2) described *in vitro* selection of TBD09-resistant mutants with inactivating mutations in *Rv3160c*, a gene encoding a TetR-family transcriptional regulator of *Rv3161c*, with a spontaneous frequency of resistance as high as 1 in 10^6^. Although we did not evaluate the bacteria isolated at Week 8 for TBD09 resistance, the lack of an observable bactericidal effect of TBD09 monotherapy at Week 4 makes it highly improbable that TBD09 monotherapy led to selective amplification of such infrequent TBD09-resistant mutants to the extent that they would overtake the TBD09-susceptible majority population in 4-8 weeks. However, as *Rv3161c* expression increases as part of the core lipid response of *M. tuberculosis* to the availability of nutrient lipid substrates *in vitro* and in mice (13, 14), the possibility that transient changes in *Rv3161c* expression over time during mouse infection contributes to some degree of tolerance to TBD09 through accelerated drug modification may be considered.

We did not evaluate safety in this study but the preclinical safety profile of TBD09 has been favorable and no dose-related trends in hematological or other safety laboratory parameters was observed over 28 days, nor were any adverse events of special interest or Grade 3 or higher adverse events observed in the phase 1 trial (2,9).

In conclusion, the results presented here suggest that TBD09 can replace the additive bactericidal and sterilizing activity of linezolid in combination with BPa and BPaM. Taken together with previously reported pre-clinical and early clinical results suggesting improved tolerability and a potentially lower risk of mitochondrial toxicity, these findings justify further clinical study of novel drug regimens based on the backbone of BPa(M) plus TBD09.

## MATERIALS AND METHODS

### Bacterial strain

*M tuberculosis* H37Rv (ATCC 27294) was used for the study. The strain was passaged in mice, confirmed to be virulent and then frozen at-80 °C. Prior to infection, one aliquot was thawed and sub-cultured in Middlebrook 7H9 broth supplemented with 10% (vol/vol) oleic acid-albumin-dextrose-catalase (OADC) enrichment (Becton-Dickinson) and 0.05% (vol/vol) Tween 80 (Fisher Scientific). This inoculum was used for aerosol infection when the optical density at 600 nm was approximately 1.0.

### Antimicrobials

Pyrazinamide was purchased from Sigma (USA), moxifloxacin from BioSynth (USA), and sutezolid from Neuland Laboratories Ltd (India). Linezolid was provided by Evotec, TBD09 by Merck (USA), bedaquiline and pretomanid by TB Alliance (New York, NY). Dosing formulations were prepared as follows: pyrazinamide and moxifloxacin in distilled water, bedaquiline in acidified 10% (2-hydroxypropyl)-ß-cyclodextrin (HPCD) solution, pretomanid in 20% HPCD and lecithin micelle (CM-2) formulation, sutezolid in a vehicle of 5% polyethylene glycol 200 (PEG 200) and 95% methylcellulose (0.5%), linezolid in 0.5% methylcellulose, and TBD09 in a vehicle of 10% Tween-80, 40% PEG 400 and 50% water. Drugs were administered orally via gavage once daily, 5 days per week in all experiments. Bedaquiline and pretomanid were administered together first, other drugs alone or in combination were administered 3 hours later.

### MIC determination

MICs were determined by the broth macrodilution method (for TBD09, linezolid and sutezolid) and by the resazurin microplate assay method (for TBD09 and linezolid). For the broth macrodilution method, *M. tuberculosis* H37Rv was grown in Middlebrook 7H9 broth supplemented with 10% OADC (BD), 0.5% glycerol, and 0.05% Tween. TBD09, linezolid and sutezolid stocks were prepared in 100% dimethylsulfoxide. Serial twofold dilutions of each stock solution were made with the same 7H9 media but without Tween 80 to yield final assay concentrations ranging from 0.06 to 4 µg/ml. Drug-free tubes served as growth controls. Each polystyrene tube containing 2.4 ml of two-fold diluted drug solution was inoculated with 100 µL of *M. tuberculosis* culture adjusted to 0.1 OD at 600 nm to give a final bacterial concentration of approximately 10^5^ CFU/ml Initial assessment was made after 1 week of incubation, followed by a final assessment at 2 weeks. The MIC was defined as the lowest concentration that inhibited visible bacterial growth after 14 days of incubation at 37°C. For the resazurin microtiter assay, *M. tuberculosis* H37Rv was grown in Middlebrook 7H9 broth supplemented with 10% OADC, 0.5% glycerol, and 0.05% Tween80. TBD09 and linezolid stocks were prepared in 100% dimethylsulfoxide. Serial twofold dilutions of each stock solution were made with the same 7H9 media but without Tween80 to yield final assay concentrations ranging from 0.015 to 8 µg/ml for TBD09, and from 0.03 to 16 µg/ml for linezolid. Drug-free wells served as growth controls, and wells without drug or *M. tuberculosis* served as contamination controls. Each test well of a 96-well polystyrene flat-bottom plate containing 100 µl of twofold diluted drug solution was inoculated with 100 µl of *M. tuberculosis* culture adjusted with 0.01 OD at 600 nm to give a final bacterial concentration of approximately 10^5^ CFU/ml. After 7 days of incubation, 30 µl of 0.02% resazurin solution prepared in sterile water was added to each well and left to incubate an additional 24 hours. Following incubation, the fluorescence signal from each well was measured using the FLUOstar OPTIMA (BMG Labtech), and the result exported for downstream analysis. Percent inhibition in each test well was calculated by subtracting the background fluorescence signal from the gross fluorescence, then dividing the fluorescence of each test well by the drug-free control well and converting to a percentage. Percent inhibition was plotted against log_10_-transformed drug concentrations and a nonlinear (sigmoidal Emax) curve with variable slope was fit using least squares. The MIC was defined as the concentration that inhibited fluorescence by 90% compared to the drug-free control wells. The experiment was performed twice, with 3 technical replicates per drug concentration.

### Infection model

All animal procedures were conducted according to relevant national and international guidelines and approved by the Johns Hopkins University Animal Care and Use Committee. Female BALB/c mice (Charles River, Wilmington, MA) aged 4 to 6 weeks were aerosol-infected using an inhalation exposure system (Glas-col, Inc., Terre Haute, IN) with approximately 4 log_10_ CFU of *M. tuberculosis* H37Rv log phase culture with OD_600_ of approximately 0.8. Mice were sacrificed for lung CFU counts one day (D-13) and two weeks (D0) after infection to determine the number of CFU implanted and the number present at the start of treatment, respectively.

### Experiment 1: To determine the dose-response and exposure-response relationships for the bactericidal activity of TBD09 alone and in combination with BPa following 4 and 8 weeks of therapy

Mice were randomized to one of eleven treatment groups (**Table S1**). The control groups included linezolid and sutezolid given alone, and the combinations BPa, BPaL and BPaU. The test combinations included BPa plus either TBD09_50_, TBD09_100_ or TBD09_200_, to determine the benefit of adding each oxazolidinone to the baseline BPa combination. Mice were treated for a total of 8 weeks. The bactericidal activity of each regimen was determined by performing CFU counts after 4 and 8 weeks of treatment.

### Pharmacokinetic analysis

small-volume in-life samples of whole blood were obtained from the cheek pouch of 3 mice in each arm after the Wednesday dose during the 2^nd^ week of treatment. Mice were sampled at 2, 6 and 24 hrs after TBD09 dosing (the 24 hr time point was obtained as a terminal bleed). Prior to analysis, 15 µL of blood was added to 30 µl of water, vortexed to promote cell lysis, and then 135 µL of acetonitrile were added for a 1:3 ratio (total dilution of sample 1:12). Drug concentrations were then determined using an LC-MS/MS method developed for this purpose. Tolbutamide was used as an internal standard. A Shiimadzu Nexera HPLC system was used with an Agilent Zorbax 5B-Aq 50 × 2.1 mm, 3.5 µm column. Mobile phase A was 0.1% formic acid in water, and mobile phase B was 0.1% formic acid in acetonitrile. Injection volume was 10 µl and the flow rate was 0.8 ml/minute. Mass spectrometry was carried out with an API 4000 instrument using Analyst 1.6.2 software. The ionization method was in ESI positive mode. The transitions (m/z) were 420.2/388.2 for TBD09 and 271.0/155/1 for tolbutamide. The lower limit of quantitation was 1 ng/ml.

### Experiment 2: To determine whether dose-optimized TBD09 has additive sterilizing activity when combined with BPa, with or without moxifloxacin

Mice were randomized to one of nine treatment groups (**Table S4**). The control groups included BPa and BPaM to test benefit of adding TBD09_200_ to these regimen backbones. BPaU, BPaL and BPaUM were included as comparator regimens. Finally, BPaMZ, the 4-month regimen evaluated in the SimpliciTB trial, was included as a clinical comparator but, unlike the SimpliciTB trial regimen, pyrazinamide was given for the first 2 months only. Lung CFU counts were determined after 2 and 4 weeks of treatment to determine the bactericidal activity of each regimen, while the sterilizing activity of drug regimens was determined by holding mice for relapse after predetermined durations of treatment ranging from 4 to 22 weeks.

### Evaluation of drug efficacy

Lung CFU counts were determined at predetermined timepoints during treatment and the presence or absence of relapse after differing durations of treatment was assessed by holding cohorts of 3-5 mice per group for an additional 12 weeks without treatment and plating the entire lung homogenate. At each time point, lungs were removed aseptically and homogenized in 2.5 ml of PBS. Lung homogenates were plated in serial dilutions on 0.4% charcoal-supplemented 7H11 agar with selective antibiotics: cycloheximide (20 µg/ml), carbenicillin (100 µg/ml), polymyxin B (400,000 U/ml), and trimethoprim (40 µg/ml). Other than the portion of the homogenate used for dilutions, the entire homogenate was plated undiluted for relapse assessment. Colonies were counted after 4 and 6 weeks of incubation at 37°C. Relapse was defined by the detection of ≥1 CFU.

## Statistical analysis

Individual mouse CFU counts (*x*) were log-transformed (as *x* + 1) before analysis. Group means were compared by one-way analysis of variance with Dunnett’s posttest to control for multiple comparisons. Group relapse proportions were compared using Fisher’s exact test, adjusting for multiple comparisons. GraphPad Prism version 9 (GraphPad, San Diego, CA) was used for all analyses.

### Statistical modeling of relapse data

Statistical modeling was performed as previously described (5, 15). Model fitting was performed in R (v. 4.3.0) (16) via RStudio Pro (v. 2023.12.1) using the software program Stan (v. 2.32.2) (17) as implemented through the rstan package (v. 2.32.6) (18). Relapse data from the present study was incorporated into a pooled dataset for a total of 30 RMM studies (5, 15). Relapse was treated as a binary 0 or 1 endpoint corresponding to absence or presence of relapse, respectively, with treatment duration as an independent variable, and study as a random effect. Separate fixed effects model parameters (T_50_ and γ) were retained for each regimen in the study except for BPaL which was combined with historical data for BPaL/BPa for model stability. The resulting fitted model parameters, including historical estimates for HRZE/HR from the pooled dataset as a comparator, were used to generate the cure probability versus treatment duration curves after adjustment for the study-specific inoculum amount as a covariate. The treatment duration to 95% cure (T_95_) values, equivalent to treatment duration to 5% relapse probability, were obtained from the estimated model parameters.

## Supplemental tables and figures

**Table S1.** Scheme for Experiment 1.

|  | Time point and number of mice |  |  |  |  |  |
| --- | --- | --- | --- | --- | --- | --- |
| Treatment<sup>1</sup> | D-14 | D0 | W2 PK<sup>2</sup> | W4 | W8 | Total |
| Untreated | 4 | 6 |  |  |  | 10 |
| L<sub>100</sub> |  |  |  | 5 | 5 | 10 |
| U<sub>50</sub> |  |  |  | 5 | 5 | 10 |
| TBD09<sub>50</sub> |  |  | 3 | 5 | 5 | 13 |
| TBD09<sub>100</sub> |  |  | 3 | 5 | 5 | 13 |
| TBD09<sub>200</sub> |  |  | 3 | 5 | 5 | 13 |
| B<sub>25</sub>Pa<sub>50</sub> |  |  | 3 | 5 | 5 | 13 |
| BPaL |  |  |  | 5 | 5 | 10 |
| BPaU |  |  |  | 5 | 5 | 10 |
| BPaTBD09<sub>50</sub> |  |  | 3 | 5 | 5 | 13 |
| BPaTBD09<sub>100</sub> |  |  | 3 | 5 | 5 | 13 |
| BPaTBD09<sub>200</sub> |  |  | 3 | 5 | 5 | 13 |
| **Total** | **4** | **6** | **21** | **55** | **55** | **141** |

<sup>1</sup> **B** = Bedaquiline; **Pa** = Pretomanid; **L** = Linezolid; **U** = Sutezolid; subscript is daily dose in mg/kg
<sup>2</sup> **W2 PK** = mice for PK sampling during the 2<sup>nd</sup> week of treatment only, not for efficacy endpoint

**Table S2.** Mean lung CFU counts from Experiment 1.

|  | **Mean (SD) lung log<sub>10</sub> *M. tuberculosis* CFU at indicated time points** |  |  |  |
| --- | --- | --- | --- | --- |
| **Treatment<sup>1</sup>** | **D-13** | **D0** | **W4** | **W8** |
| Untreated | 4.46 ± 0.11 | 7.63 ± 0.06 |  |  |
| L<sub>100</sub> |  |  | 7.52 ± 0.06 | 6.28 ± 0.07 |
| U<sub>50</sub> |  |  | 6.66 ± 0.12 | 5.11 ± 0.15 |
| TBD09<sub>50</sub> |  |  | 7.44 ± 0.16 | 7.06 ± 0.19 |
| TBD09<sub>100</sub> |  |  | 7.55 ± 0.03 | 7.17 ± 0.07 |
| TBD09<sub>200</sub> |  |  | 7.65 ± 0.07<sup>1</sup> | 7.25 ± 0.11 |
| B<sub>25</sub>Pa<sub>50</sub> |  |  | 6.47 ± 0.16 | 4.37 ± 0.20<sup>1</sup> |
| BP<sub>a</sub>L |  |  | 5.36 ± 0.17 | 2.84 ± 0.16 |
| BP<sub>a</sub>U |  |  | 3.91 ± 0.33 | 1.52 ± 0.64 |
| BP<sub>a</sub>TBD09<sub>50</sub> |  |  | 5.44 ± 0.10 | 3.46 ± 0.40 |
| BP<sub>a</sub>TBD09<sub>100</sub> |  |  | 5.37 ± 0.22 | 3.15 ± 0.24 |
| BP<sub>a</sub>TBD09<sub>200</sub> |  |  | 5.49 ± 0.14 | 2.15 ± 0.26 |
| <sup>1</sup>mean CFU count from 4 mice; homogenizer broke while processing lungs, causing sample loss for one mouse in each indicated group |  |  |  |  |

**Table S3.**
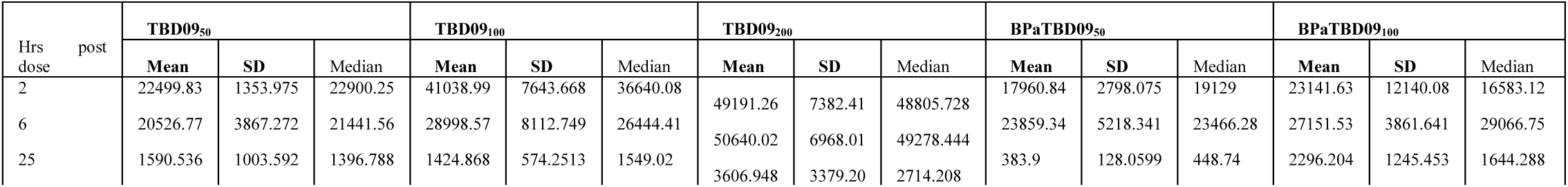
Mean and median TBD09 whole blood concentrations from Experiment 1.

**Table S4.** Scheme for Experiment 2.

| Treatment<sup>1</sup> | Time point and number of mice<sup>2</sup> |  |  |  |  |  |  |  |  |  |  |  | Total |
| --- | --- | --- | --- | --- | --- | --- | --- | --- | --- | --- | --- | --- | --- |
|  | D-13 | D0 | W2 | W4<br>(+12) | W6<br>(+12) | W8<br>(+12) | W10<br>(+12) | W12<br>(+12) | W14<br>(+12) | W16<br>(+12) | W19<br>(+12) | W22<br>(+12) |  |
| Untreated | 6 | 9 |  |  |  |  |  |  |  |  |  |  | 15 |
| B<sub>25</sub>Pa<sub>50</sub> |  |  | 5 | 5 |  |  | (4) | (5) | (5) | (5) | (4) | (3) | 10 (26) |
| BP<u>a</u>U<sub>50</sub> |  |  | 5 | 5 | (4) | (5) | (5) | (5) | (4) | (3) |  |  | 10 (26) |
| BP<u>a</u>L<sub>100</sub> |  |  | 5 | 5 | (4) | (5) | (5) | (5) | (5) | (3) | (3) |  | 10 (30) |
| BP<u>a</u>TBD09<sub>200</sub> |  |  | 5 | 5 | (4) | (5) | (5) | (5) | (5) | (3) | (3) |  | 10 (30) |
| BP<u>a</u>M<sub>100</sub> |  |  | 5 | 5 | (4) | (5) | (5) | (5) | (5) | (3) | (3) |  | 10 (30) |
| 2BP<u>a</u>MZ<sub>150</sub>/<br>BP<u>a</u>M |  |  | 5 | 5 (4) | (5) | (5) | (4) |  |  |  |  |  | 10 (18) |
| BP<u>a</u>MU |  |  | 5 | 5 (3) | (5) | (5) | (5) | (5) | (3) |  |  |  | 10 (26) |
| BP<u>a</u>MTBD09 |  |  | 5 | 5 | (5) | (5) | (5) | (5) | (5) | (5) |  |  | 10 (30) |
| **Total (n=311)** | **6** | **9** | **40** | **40 (7)** | **(31)** | **(35)** | **(38)** | **(35)** | **(32)** | **(22)** | **(13)** | **(3)** | **95**<br>**(216)** |

<sup>1</sup> B = Bedaquiline; Pa = Pretomanid; L = Linezolid; U = Sutezolid; M=Moxifloxacin; Z=pyrazinamide; subscript is daily dose in mg/kg
<sup>2</sup> Numbers outside parentheses indicate mice for lung CFU counts; numbers inside parentheses indicate mice held for 12 wks after treatment completion (+12) for relapse assessment.

**Table S5.** Mean lung CFU counts from Experiment 2.

|  | Time point and number of mice |  |  |  |
| --- | --- | --- | --- | --- |
| Treatment<sup>1</sup> | D-13<br>2/28 | D0<br>3/14 | W2 | W4 |
| Untreated | 4.41±0.14 | 7.78±0.15 |  |  |
| B<sub>25</sub>Pa<sub>50</sub> |  |  | 7.20± 0.18 | 6.09± 0.23 |
| BPaU<sub>50</sub> |  |  | 6.17 ± 0.26 | 3.66± 0.43 |
| BPaL<sub>100</sub> |  |  | 6.79 ± 0.07 | 4.75± 0.21 |
| BPaTBD09<sub>200</sub> |  |  | 6.77 ±0.08 | 4.82± 0.28 |
| BPaM<sub>100</sub> |  |  | 6.86 ±0.27 | 5.38± 0.23 |
| 2BPaMZ<sub>150</sub>/BPaM |  |  | 5.77±0.09 | 2.60± 0.34 |
| 2BPaUM |  |  | 5.97±0.16 | 3.94± 0.18 |
| BPaMTBD09 |  |  | 6.31±0.20 | 4.64± 0.14 |

<sup>1</sup>B = Bedaquiline; Pa = Pretomanid; L = Linezolid; U = Sutezolid; M=Moxifloxacin; Z=pyrazinamide; subscript is daily dose in mg/kg

**Table S6.** Treatment duration to 95% cure (T95) values from model-based relapse analysis.

| Regimen | T95 Metrics |  |  |  |  |
| --- | --- | --- | --- | --- | --- |
|  | Mean | SD | 5th | Median | 95th |
| BPaMZ | 1.034 | 0.1902 | 0.745 | 1.016 | 1.368 |
| BPaM + TBD09 | 2.715 | 0.4839 | 2.159 | 2.636 | 3.517 |
| BPa + TBD09 | 2.717 | 0.2940 | 2.322 | 2.679 | 3.230 |
| BPaL | 3.209 | 0.1186 | 3.021 | 3.203 | 3.407 |
| BPaM | 3.284 | 0.4247 | 2.759 | 3.216 | 4.035 |
| BPaU | 3.421 | 3.3090 | 1.899 | 2.816 | 6.269 |
| BPaMU | 3.710 | 4.3020 | 2.286 | 3.089 | 6.058 |
| BPa | 4.269 | 0.1263 | 4.064 | 4.264 | 4.486 |
| HRZE/HR | 5.604 | 0.1833 | 5.324 | 5.594 | 5.922 |

---

**Figure S1.**
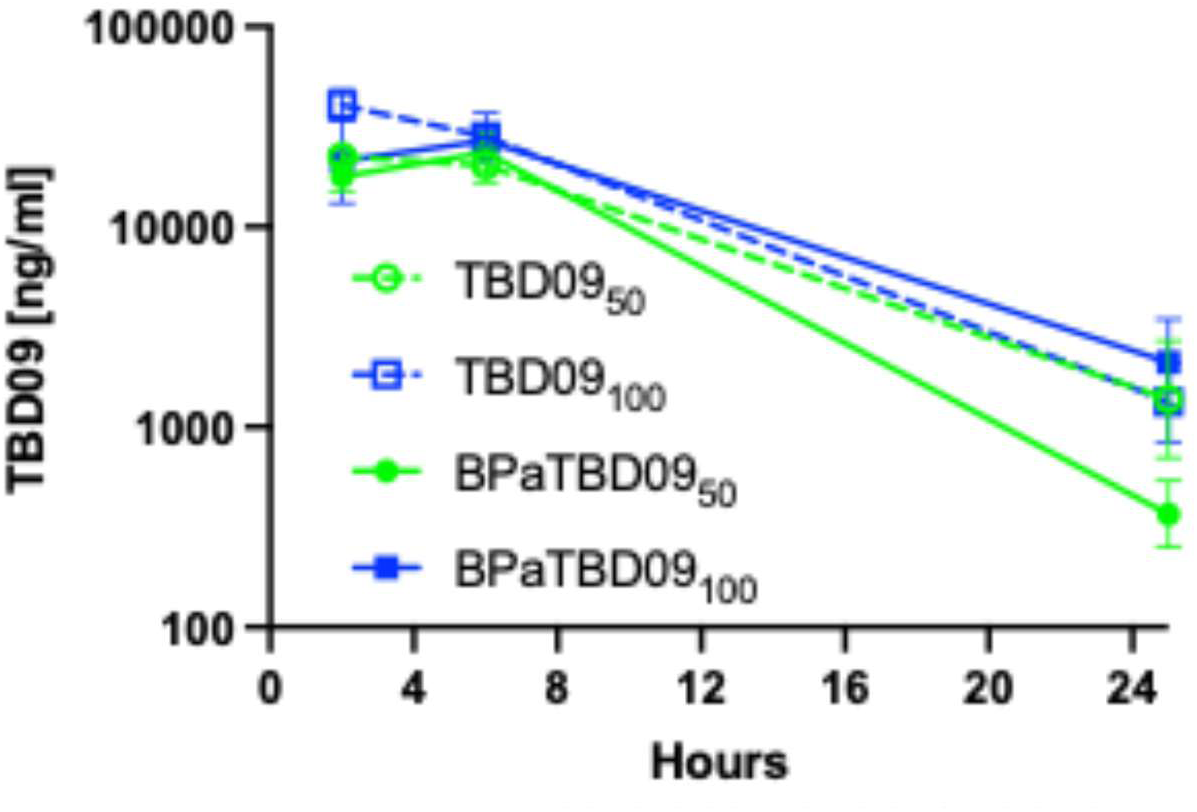
Mean TBD09 whole blood concentrations from Experiment 1

**Figure S2.**
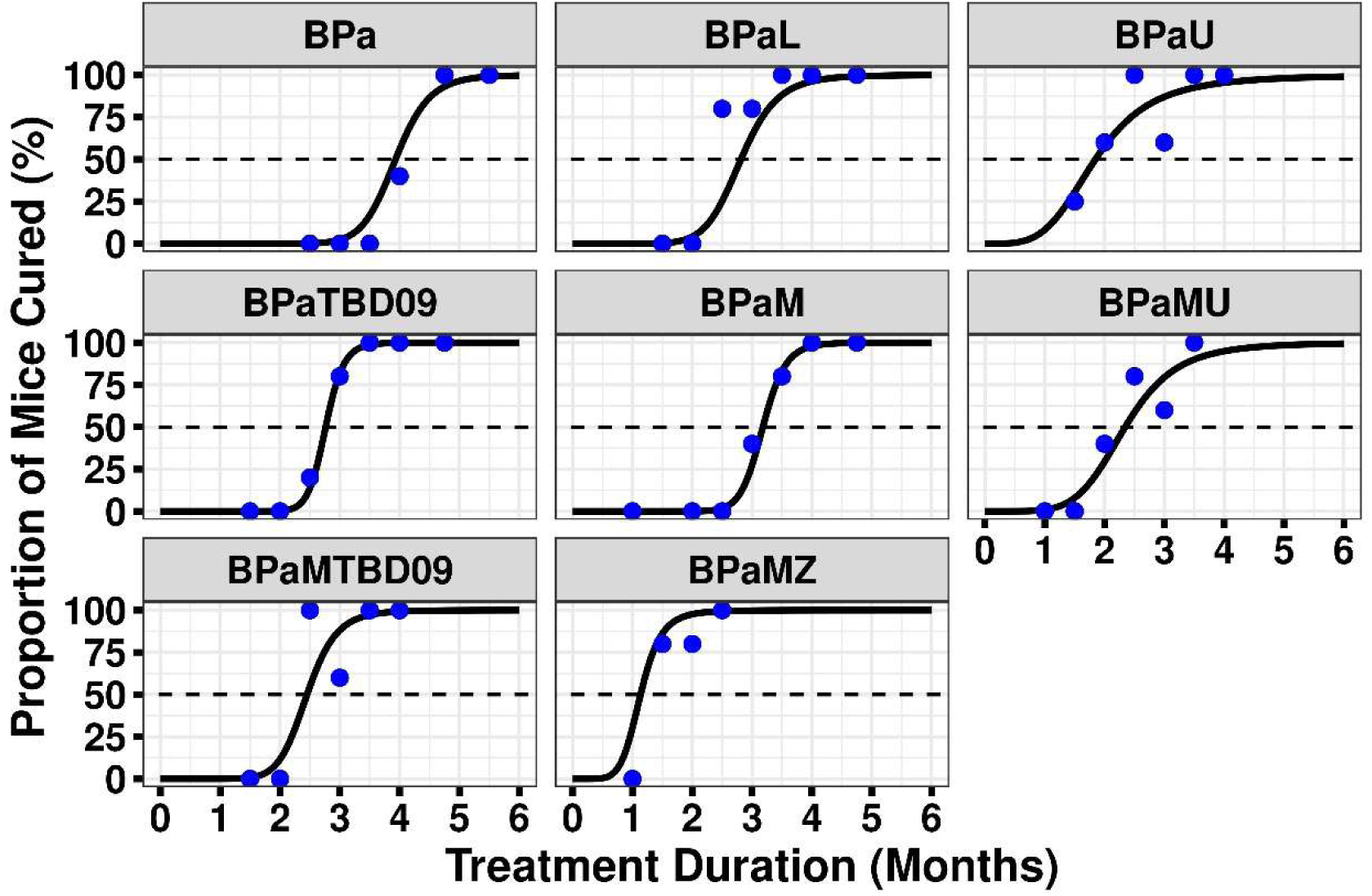
Proportion of mice cured by duration of treatment for each study arm

## REFERENCES

1. World Health Organization. 2022. WHO Guidelines Approved by the Guidelines Review Committee, WHO consolidated guidelines on tuberculosis: Module 4: treatment - drug-resistant tuberculosis treatment, 2022 update. World Health Organization.

2. Crowley BM, Boshoff HI, Boving A, Tan VY, Zhu J, Hoyt F, Miller RR, Ehrhart J, Boyce CW, Young K, Nantermet PG, Su J, Yang L, Painter RE, Corcoran EB, Hoar JL, Oh S, Holtzman DL, Levi M, Anderson A, Otieno MA, Zimmerman M, Kaya F, Massoudi LM, Ramey ME, Bauman AA, Lenaerts AJ, Roberston GT, Dartois V, Wells CD, Barry CE 3rd, Olsen DB. 2026. Discovery and development of a new oxazolidinone with reduced toxicity for the treatment of tuberculosis. Nat Med 32:553–560. doi: 10.1038/s41591-025-04164-x. PMID: 41530381; PMCID: PMC12920096.

3. Tasneen R, Betoudji F, Tyagi S, Li SY, Williams K, Converse PJ, Dartois V, Yang T, Mendel CM, Mdluli KE, Nuermberger EL. 2016. Contribution of oxazolidinones to the efficacy of novel regimens containing bedaquiline and pretomanid in a mouse model of tuberculosis. Antimicrob Agents Chemother 60: 270–7.

4. Li SY, Tasneen R, Tyagi S, Soni H, Converse PJ, Mdluli K, Nuermberger EL. 2017. Bactericidal and sterilizing activity of a novel regimen with bedaquiline, pretomanid, moxifloxacin, and pyrazinamide in a murine model of tuberculosis. Antimicrob Agents Chemother 61: e00913–17. doi: 10.1128/AAC.00913-17. PMID: 28630203; PMCID: PMC5571308.

5. Berg A, Clary J, Hanna D, Nuermberger E, Lenaerts A, Ammerman N, Ramey M, Hartley D, Hermann D. 2022. Model-Based meta-analysis of relapsing mouse model studies from the critical path to tuberculosis drug regimens initiative database. Antimicrob Agents Chemother 66: e01793–21. 10.1128/aac.01793-21

6. Nunn AJ, Phillips PPJ, Meredith SK, Chiang CY, Conradie F, Dalai D, van Deun A, Dat PT, Lan N, Master I, Mebrahtu T, Meressa D, Moodliar R, Ngubane N, Sanders K, Squire SB, Torrea G, Tsogt B, Rusen ID; STREAM Study Collaborators. 2019. A trial of a shorter regimen for rifampin-resistant tuberculosis. N Engl J Med 380: 1201–1213. doi: 10.1056/NEJMoa1811867. Epub 2019 Mar 13. PMID: 30865791.

7. Sotgiu G, Centis R, D’Ambrosio L, Alffenaar J-WC, Anger HA, Caminero JA, Castiglia P, De Lorenzo S, Ferrara G, Koh W-J, Schecter GF, Shim TS, Singla R, Skrahina A, Spanevello A, Udwadia ZF, Villar M, Zampogna E, Zellweger J-P, Zumla A, Migliori GB. 2012. Efficacy, safety and tolerability of linezolid containing regimens in treating MDR-TB and XDR-TB: systematic review and meta-analysis. Eur Respir J 40:1430– 1442.

8. Kodama T, Sasaki Y, Tsuyuguchi K, Okumura M, Kamada K, Yamane A, Hayashi Y, Hagiwara E, Wakamatsu K, Kato T, Koreeda Y, Kuwabara K, Amishima M, Tamaki S, Takaki A, Kato S, Nagai H, Mitarai S, Yoshiyama T. 2025. Clinical outcomes of linezolid-related adverse events in patients with multidrug-resistant TB. Int J Tuberc Lung Dis 29:462–468. doi: 10.5588/ijtld.25.0125. PMID: 41410988.

9. Holtzman D, Vinnard C, Anderson A, Yeakey A, Daglish L, McIntyre E, Nussbaum J, Stamm L, Alexander J, Rasmussen S, Levi M, Wells C. Abstr. 56^th^ Union World Conference on Lung Health, Copenhagen November 19, 2025. LB02-1315-19 In pursuit of an improved oxazolidinone for TB: A phase 1 trial evaluating safety, tolerability, pharmacokinetics, and food effect of TBD09 (MK-7762) in healthy adults.

10. Diacon AH, De Jager VR, Dawson R, Narunsky K, Vanker N, Burger DA, Everitt D, Pappas F, Nedelman J, Mendel CM. 2020. Fourteen-Day bactericidal activity, safety, and pharmacokinetics of linezolid in adults with drug-sensitive pulmonary tuberculosis. Antimicrob Agents Chemother 64: e02012–19. doi: 10.1128/AAC.02012-19. PMID: 31988102; PMCID: PMC7179319.

11. Imperial MZ, Nedelman JR, Conradie F, Savic RM. 2022. Proposed linezolid dosing strategies to minimize adverse events for treatment of extensively drug-resistant tuberculosis. Clin Infect Dis 74: 1736–1747. doi: 10.1093/cid/ciab699. PMID: 34604901; PMCID: PMC9155613.

12. Pi R, Liu Q, Jiang Q, Gao Q. 2019. Characterization of linezolid-resistance-associated mutations in Mycobacterium tuberculosis through WGS. J Antimicrob Chemother 74: 1795–1798. doi: 10.1093/jac/dkz150. PMID: 31225608.

13. Aguilar-Ayala DA, Tilleman L, Van Nieuwerburgh F, Deforce D, Palomino JC, Vandamme P, Gonzalez-Y-Merchand JA, Martin A. 2017. The transcriptome of Mycobacterium tuberculosis in a lipid-rich dormancy model through RNAseq analysis. Sci Rep. 7:17665. doi: 10.1038/s41598-017-17751-x. PMID: 29247215; PMCID: PMC5732278.

14. Lawrence AE, Tan S. 2026. Direct bacterial transcript visualization illuminates spatiotemporal environmental adaptation of Mycobacterium tuberculosis in the lung. Sci Adv. 12: eaeb8377. doi: 10.1126/sciadv.aeb8377. Epub 2026 Aug 12. PMID: 42585320; PMCID: PMC13464648.

15. Clary J, Roberts JK, Hanna D, Tagliavini A, Sordello S, Upton A, Hermann D, Berg A. 2026. A stochastic simulation-based approach to inform the relapsing mouse model study design for non-clinical assessment of tuberculosis. Antimicrob Agents Chemother 70: e0110325. doi: 10.1128/aac.01103-25. Epub 2025 Dec 29. PMID: 41459930; PMCID: PMC12888889.

16. R Core Team (2023). R: A language and environment for statistical computing. R foundation for Statistical Computing, Vienna, Austria. https://www.R-project.org/

17. Stan Development Team. 2023. Stan Modeling Language Users Guide and Reference Manual, 2.32.2. https://mc-stan.org/docs/2_32/reference-manual/

18. Stan Development Team (2024). “RStan: the R interface to Stan.” R package version 2.32.6, https://mc-stan.org/.

